# A subcellular cookie cutter for spatial genomics in human tissue

**DOI:** 10.1101/2021.11.29.470247

**Authors:** Alexander G. Bury, Angela Pyle, Fabio Marcuccio, Doug M. Turnbull, Amy E. Vincent, Gavin Hudson, Paolo Actis

## Abstract

Intracellular heterogeneity contributes significantly to cellular physiology and, in a number of debilitating diseases, cellular pathophysiology. This is greatly influenced by distinct organelle populations and to understand the aetiology of disease it is important to have tools able to isolate and differentially analyse organelles from precise location within tissues. Here we report the development of a subcellular biopsy technology that facilitates the isolation of organelles, such as mitochondria, from human tissue. We compared the subcellular biopsy technology to laser capture microdissection (LCM) that is the state of art technique for the isolation of cells from their surrounding tissues. We demonstrate an operational limit of (>20μm) for LCM and then, for the first time in human tissue, show that subcellular biopsy can be used to isolate mitochondria beyond this limit.

## Introduction

Inter-tissue and inter-cellular heterogeneity is a known contributor to a number of human diseases including cancer [1–3]; cardiovascular disease [4,5]; metabolic disease [6–9]; neurodegeneration, neurodevelopmental disorders and pathological ageing [10–13]. Yet, evaluating heterogeneity at the tissue and cellular level can often mask subtle subcellular and organelle heterogeneity [14]. In addition to morphological and functional heterogeneity, exhibited by other organelles, mitochondria additionally show genetic heterogeneity, owing to their own multi-copy genome [9,15]. The mitochondrial genome (mtDNA) exists as uniform wild-type molecules at birth, in healthy individuals - termed homoplasmy, but *de novo* mutations give rise to a mixture of wild-type and mutant mtDNA molecules – termed heteroplasmy [16]. Whilst low level heteroplasmy is well tolerated, the accumulation and spread of mutant mtDNA molecules in excess of a threshold level, can lead to impaired oxidative phosphorylation that often culminates in mitochondrial disease [16,17]. The mechanism behind this process, termed clonal expansion, is not fully understood. Investigating clonal expansion at the subcellular level may advance our understanding of the mechanisms behind it and help improve characterisation of mitochondrial disease [18,19]. More generally, better understanding the physiological (and pathophysiological) relevance of intracellular organelle heterogeneity with subcellular precision would likely aid effective diagnsis and treatment of disease, however, we need the appropriate technologies to achieve this [9,20,21].

To take full advantage of ‘single-cell multiomics’ [22, 23], nanoprobe-based technologies can circumvent common challenges associated with investigating subcellular molecules, including: the requirement to permeabilize cells and implementation of complex biochemical reactions which do not allow, for example, the analysis of intracellular organelles [24–28]. Nanoprobe technologies are typically integrated with scanning probe microscopy to enable their automated positioning in and around cells with nanometer precision [29,30]. Hence, the comparative small probe size enables sampling from live cells with minimal impact on cell viability or the cellular environment [31].

In 2014, Actis and colleagues developed nanobiopsy technology which utilises a nanopipette containing an organic solvent to aspirate mRNA and mitochondria within the cytoplasm of cultured fibroblasts [26]. This methodology relies on electrowetting, a process where a liquid-liquid interface is manipulated by the application of a voltage to aspirate a target from the cytoplasm of a living cell [29–31]. More recently, fluid force microscopy (FFM), dielectrophoretic nanotweezers and nanopipettes have successfully been used to sample cytoplasmic proteins and nucleic acids [26,32–36] from cells in culture. However, none of these technologies have been applied to the study of tissue samples that are routinely used for clinical and molecular pathology [8,37,38]. Here, we aimed to determine if nanobiopsy could be adapted for sampling from human tissue samples.

Electrowetting is necessary for the sampling from cultured cells because manipulating the liquid-liquid interface creates a force that draws the cytoplasm into the pipette tip [31]. Our data shows that the nanopipette effectively acts as a subcellular “cookie cutter” removing the need for electrowetting (Figure 1). This is important because electrowetting relies on the use of a toxic organic solvent, 1,2-dichloroethane, that can potentially affect the quality of downstream molecular analyses [39]. Using laser-capture-microdissection (LCM), the most commonly utilised approach for studying single cells from tissue samples [10,18,38,40,41], as a comparator, we show that an adaptation of nanobiopsy: subcellular biopsy, has the potential to go beyond conventional methodological limits.

**Figure 1.**
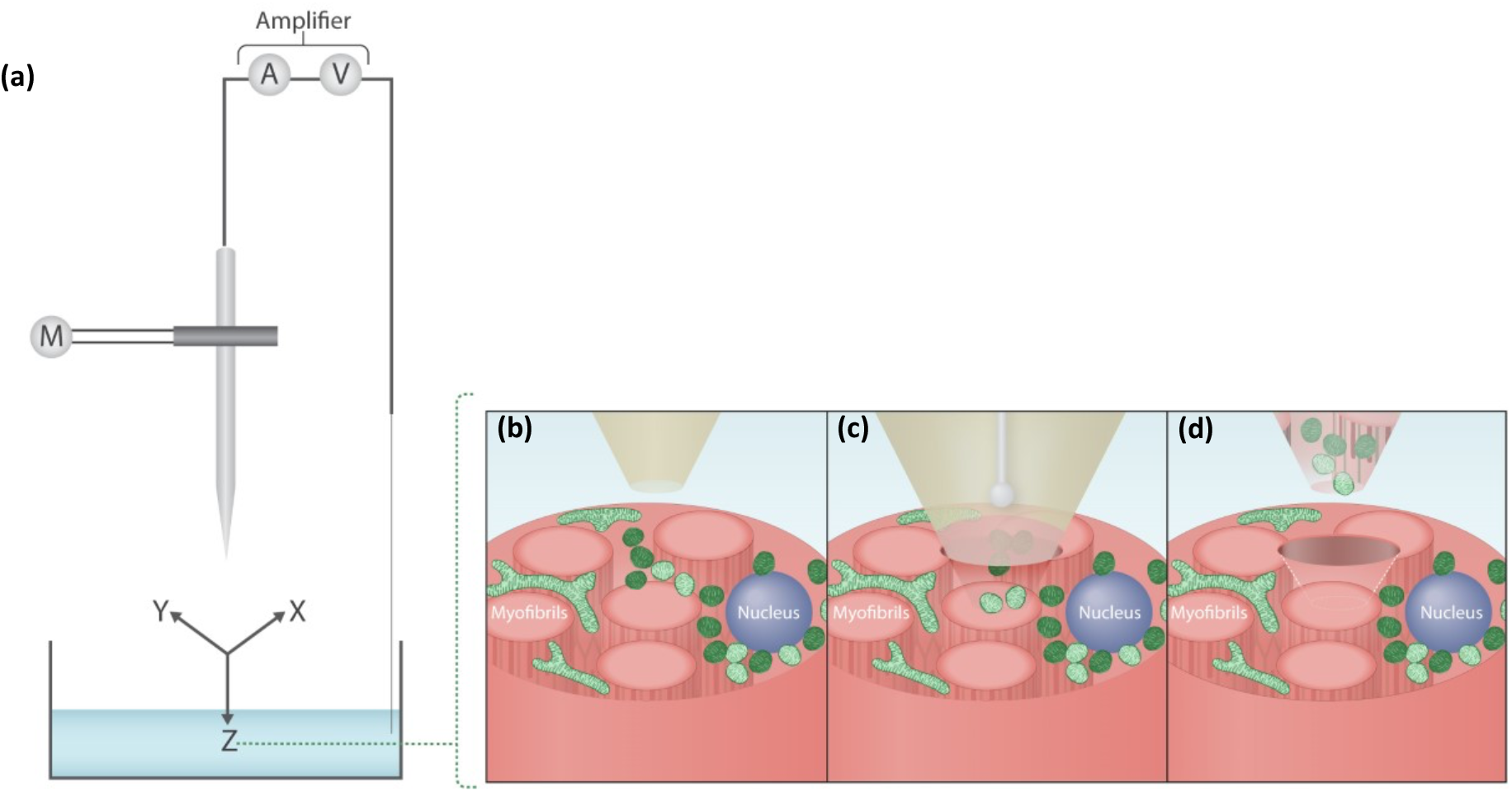
(a) Subcellular biopsy in skeletal muscle. A micropipette is incorporated within a scanning ion conductance microscope to enable its automated positioning **(b)** A micropipette is immersed within an aqueous bath in which the sample of interest is placed, such as skeletal muscle tissue. The micropipette can be positioned within 1μm of the skeletal muscle fibre of interest **(c)** The micropipette then penetrates the tissue by a predefined distance to perform the subcellular biopsy **(d)** The micropipette is then retracted and the biopsied content are placed into a collection vessel.

## Materials and Methods

### Skeletal Muscle Tissue Biopsies

Skeletal muscle tissue biopsies (*n* = 14) were obtained from volunteers undergoing anterior cruciate ligament (ACL) surgery at the Royal Victoria Infirmary (Newcastle, UK). Consent and ethical approval were granted by The Newcastle and North Tyneside Local Research Ethics Committee (LREC: 12/NE/0395). All research was carried out in correspondence with the human tissue act (HTA) (2004) with HTA licensing and material transfer agreements between the Newcastle University and University of Leeds. Biopsied tissue was snap frozen in liquid nitrogen cooled isopentane and mounted transversely in OCT mounting medium. Cryosections were collected onto slides (Agar Scientific Ltd, Essex, UK), air-dried for 1 hour and stored at −80°C prior to staining.

### Immunofluorescence Staining

Briefly, tissue sections were fixed with 4% PFA (Sigma-Aldrich, Missouri, US) and permeabilised using a methanol gradient (ThermoFisher, Paisley, UK). A protein block (10% normal goat serum, Vector, Peterborough, UK) was carried out at room temperature for 1 hour and endogenous biotin blocking (Avidin D/Biotin, Vector) was carried out applying each solution for 15 mins at room temperature, to minimise non-specific staining. Primary and secondary antibodies were applied overnight and for 2 hours respectively, at 4°C in a humidified chamber. Primary antibodies: anti-mitochondrial cytochrome c oxidase I (MTCOI; ab14705, Abcam, Cambridge, UK), anti-succinate dehydrogenase subunit A (SDHA; ab14715, Abcam); anti-Voltage Dependent Anion Channel 1 (VDAC1; ab14734, Abcam); 4,6-diamidino-2-phenylindole nuclear stain (DAPI; Sigma, St Louis, MO). Secondary antibodies (all Life Technologies): anti-mouse IgG2a-488 (S21131), biotinylated anti-mouse IgG1 (S32357), streptavidin-647 (S21374) anti-mouse IgG2b-546 (L9168).

### Laser Capture Microdissection (LCM) and Tissue Lysis

Isolation of single- and subcellular muscle fibre dissections were performed using a PALM LCM system (Zeiss, Oberkochen, DE). Microdissected samples were collected in inverted 0.2ml microfuge tubes containing 15μl of lysis buffer (0.5M Tris-HCl, 0.5% Tween 20, 1% proteinase K at pH 8.5), as previously described [41]. Samples were centrifuged and then incubated at 55°C for 3 hours then at 95°C for 10 mins, using a thermal cycler (Applied Biosystems), to ensure efficient lysis of single cells and mitochondria to yield mtDNA.

### Fabrication and analysis of micropipettes

Micropipettes were fabricated from quartz glass capillaries (QF100-50-7.5, Sutter Instrument, Novato, CA) using a P-2000 micropipette laser puller (World Precision Instruments, Sarasota, FL). An Ag/AgCl wire was inserted in the micropipette to act as the working electrode and another Ag/AgCl wire was immersed in a 1xPBS bath (Oxoid Ltd, ThermoFisher, UK), acting as the reference electrode. Current-voltage measurements are performed using a MultiClamp 700B patch-clamp amplifier (Molecular Devices, Sunnyvale, CA). The signal was filtered using a Digidata 1550B digitizer, with a low-pass filter at 10kHz, and signal recording was performed using the pClamp 10 software (Molecular Devices), at a rate of 100kHz. For electrochemical analysis experiments, the micropipettes were filled with 1xPBS. For electrowetting experiments, the micropipettes were filled with a solution of 10mM tetrahexylammonium tetrakis(4-chlorophenyl)borate (THATPBCl) salt in 1,2-dichloroethane (1,2-DCE).

### Scanning Electron Microscopy (SEM)

A Vega 3 Scanning Electron Microscope (Tescan, Brno, CZ) was used to image and determine the micropipette geometry and aperture size. Micropipettes were sputter coated with a 5nm layer of gold before being mounted onto a sample holder. Imagine parameters were as follows: acceleration voltage: 8kV; beam intensity: 6-8; working distance: 15-30mm; magnification: X10k-60k.

### Scanning Ion-Conductance Microscopy (SICM)

The SICM set-up was comprised of an Axon MultiClamp 700B amplifier, MM-6 micropositioner (Sutter Instrument, Novato, CA) and a P-753 Linear actuator (Physik Instrumente, Irvine, CA) attached to the pipette holder to allow precise, three-dimensional movement of the micropipette (Figure 1a). SICM software was used to control the positioning and topographical scanning capabilities of the SICM system (ICAPPIC, London, UK). An Eclipse Ti2 confocal microscope (Nikon Instruments Inc., Melville, NY) and broad-spectrum LED illumination system (pE-300 CoolLED, Andover, US) were used for bright-field (BF) and immunofluorescence (IF) visualisation of mitochondria and myonuclei, in skeletal muscle fibres, to ensure efficient lysis. The SICM system is used to automatically approach the skeletal muscle tissue, moving the micropipette just above a region of interest (Figure 1b) and then to enter the skeletal muscle fibre through manual control (Figure 1c). Following successfully biopsy of mitochondria, the micropipette was retracted the tip snapped into a 0.2ml microfuge tubes containing lysis buffer (Figure 1d): *as described in the LCM section above*.

### Triplex Quantitative PCR (qPCR)

To quantify total mtDNA CN the abundance of the *MTND1* probe (mtDNA minor arc) was measured, indicative of total mtDNA [42]. The *D Loop* probe, corresponding to another highly conserved region of the mitochondrial genome, was used as a comparator in case of rare instances of minor-arc deletions that could affect accurate CN assessment [43]. Samples were plated in triplicate and each qPCR run was repeated, to control for inter-run variation.

### Statistical Analysis

The variance in the median between electrowetting and non-electrowetting groups, and each of electrowetting and non-electrowetting against non-biopsy controls (NBC) were compared using Mann-Whitney U tests. The correlation between mtDNA CN and LCM dissection or subcellular biopsy size was tested by linear regression. Statistical significance was set as *p*<0.05. Statistical analyses were performed and graphs produced using GraphPad Prism version 5.00 for Windows (GraphPad Software, San Diego, CA).

## Results

### Working range of Laser Capture Microdissection (LCM)

LCM is the most established technique for isolating regions of interest within tissue samples [18,38,40]. First, optimisation of the immunostaining of 10μm skeletal muscle sections was performed using a protocol adapted from Rocha and colleagues [44], to allow visualisation of myonuclei and mitochondria. We then established the working limit of LCM, by determining the smallest possible dissected area that could then be successfully analysed using a qPCR assay targeting mtDNA. We stained a human skeletal muscle tissue slice with a fluorescently labelled antibody targeting mitochondrially encoded cytochrome c oxidase I (MTCOI) and used fluorescent microscopy to confirm the success of the staining (Figure 2ai and bi). We used LCM to dissect regions with a decreasing area and Figure 2aii shows a BF micrograph after we performed four dissections of 30 μm in diameter. Fluorescence micrographs (Figure 2aiii) also confirmed the success of the dissections. Similarly, we performed dissections of 5 μm in diameter and imaged the sample before (Figure 2bi) and after the procedure (Figure 2bii and 2biii).

**Figure 2.**
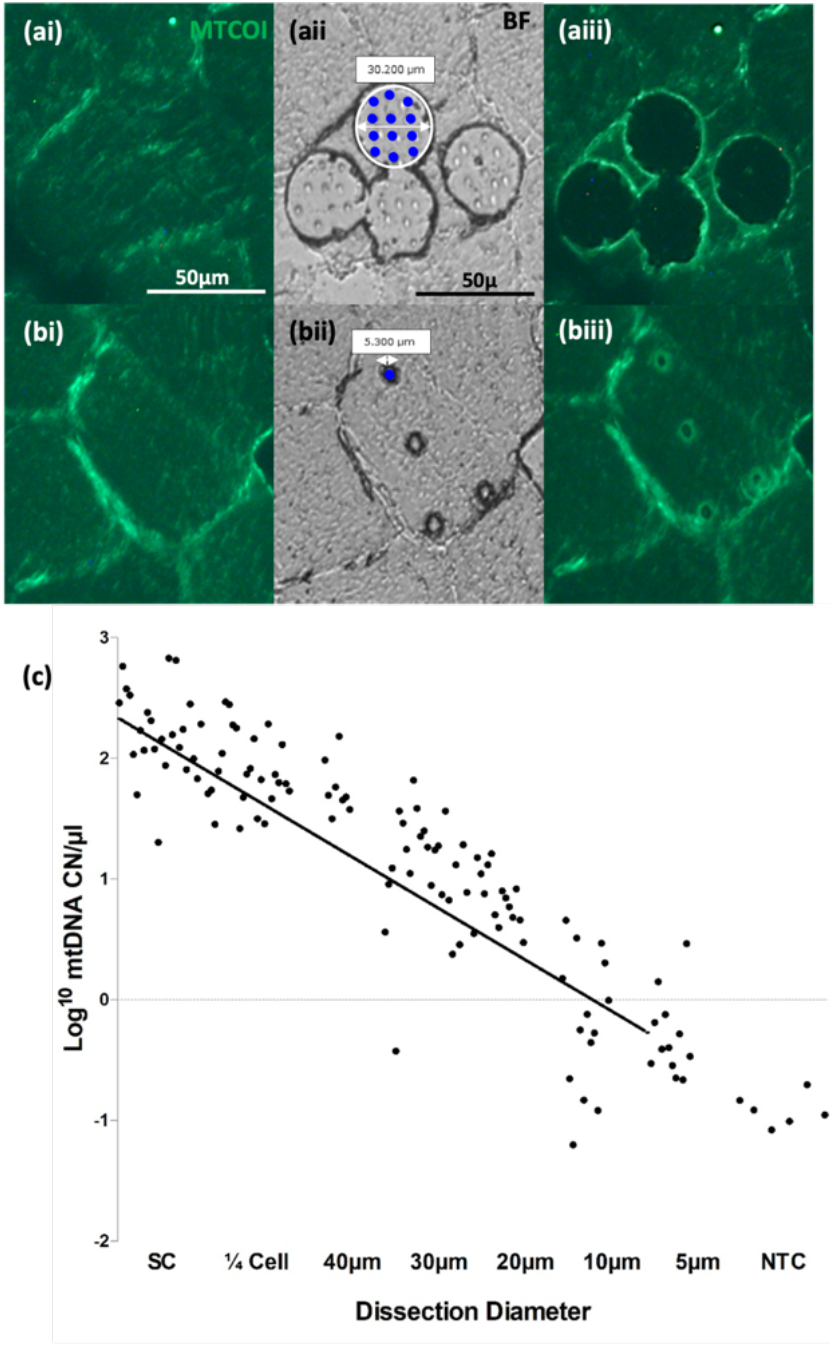
Working limit of LCM. Panels **(a)** and **(b)** correspond to biopsies taken using LCM dissection diameters of 30μm **(a)** and 5μm **(b)** in diameter. The first two panels (left to right) correspond to pre-biopsy immunofluorescent (IF) images to identify the area of interest **(ai, bi)**. The middle panels show the BF post-cut images with the delineated cutting area outlined in white **(aii)** and laser pulse markers, shown as blue dots **(aii, bii)**. The right panels correspond to post-dissection IF images showing the indentations made by the cutting laser and laser pulses and the relative change in fluorescence indicating successful isolation of tissue **(aiii, biii)**. CN/μl values (log transformed, y-axis) were plotted against LCM dissection size (μm). Bars on the graph represent 95% CI for each dissection size. Regression analysis demonstrated a significant relationship between LCM dissection size and (*n* = 14; Spearman correlation=0.91, *p<* 0.05). A greater proportion of dissections under 20μm in diameter corresponded to less than 1 mtDNA copies (< 0 log transformed CN/μl), compared with dissections larger than 20μm **(c)**.

**Figure 3.**
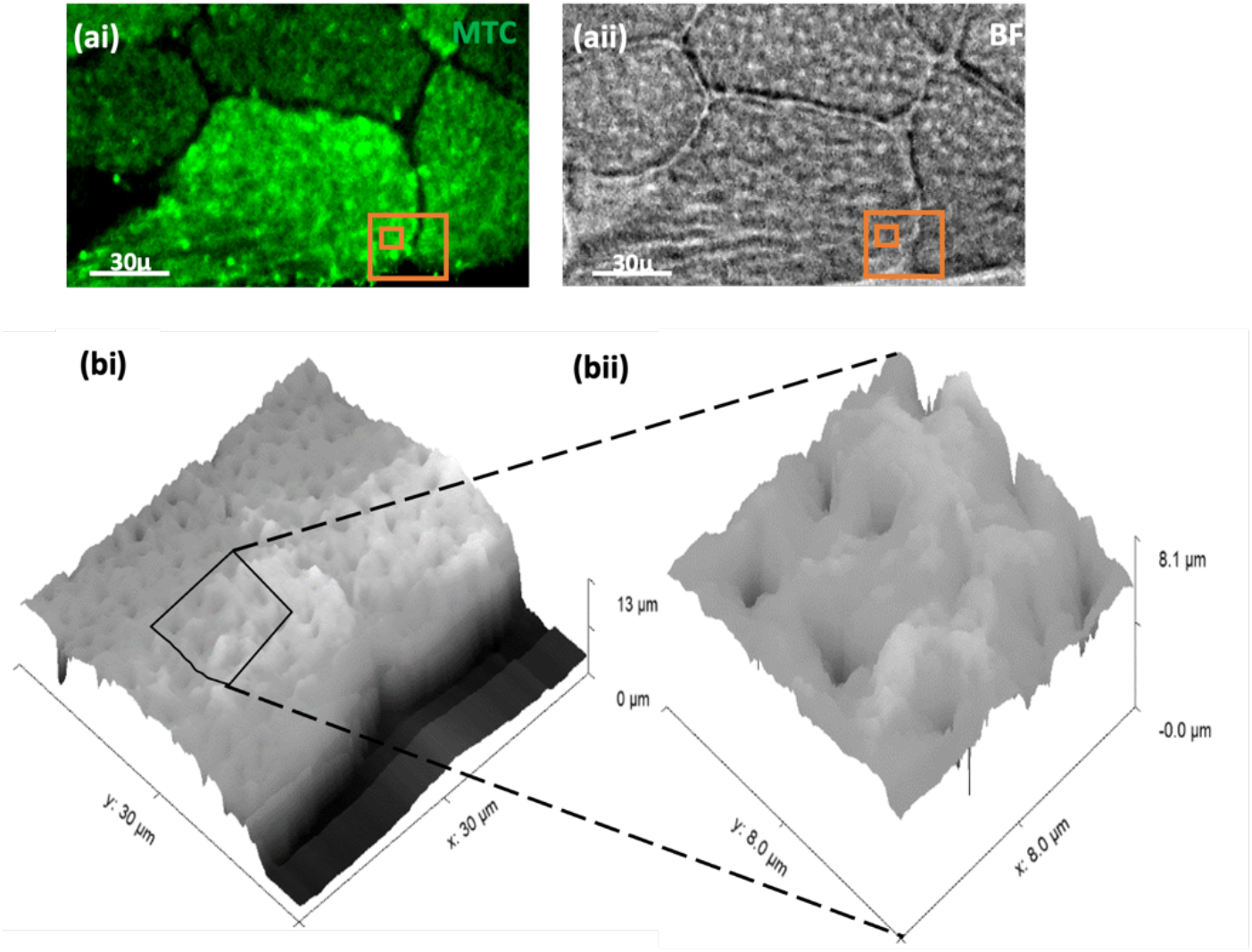
Fluorescent micrograph (ai) corresponding to IF labelling of MTCOI **(ai)** and bright field micrograph **(aii)** of a human skeletal tissue slice, with orange boxes delineating the SICM scan areas. SICM topography scans of a 30×30 μm **(bi)** and 8×8 μm **(bii)** region of interest.

A Triplex qPCR protocol was adapted, for use with subcellular quantities of mtDNA, from Rygiel and colleagues [42. Using a series of increasingly smaller diameter dissections (n=4 per section) ranging from whole single cells to 5μm dissections, we were able to show that the reliable working limit of LCM is between 20 and 10μm (Figure 2c) and we observed that the mtDNA copy number per microliter (CN/μl) is correlated to the dissected area (r = −0.91, p < 0.0001, Figure 2c). However, beyond 20 μm mtDNA copy-number assessments from repeat sampling showed increased variability and often <1 copy of mtDNA. Having demonstrated the working limit of LCM, we then investigated if an adaption of the nanobiopsy technology based on a scanning ion conductance microscope (SICM) could be implemented to enable sub cellular sampling from human tissue sections with a spatial resolution surpassing the limits of LCM.

SICM enables the topographically mapping of cells and tissues with nanoscale resolution [45]. The SICM setup is mounted on top of an inverted optical microscope to enable simultaneous optical imaging and topographic mapping of the sample of interest. Figure 2 shows the optical (panel a) imaging of a skeletal muscle tissue and its topography (panel b) as acquired with the SICM showing the sarcoplasm.

### Subcellular mitochondrial biopsy from skeletal muscle fibres

The subcellular biopsy approach relies onto a micropipette integrated into a SICM setup. Two twin micropipettes were produced from each capillary tube to specifications adapted from the accompanying manual (Sutter Instrument). An optimised two-line patch pipette programme was used, as follows:

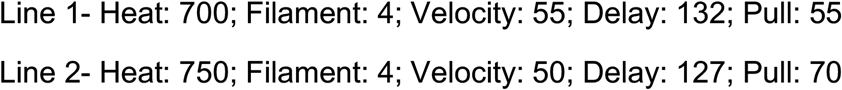

Scanning Electron Microscopy (SEM) and electrochemical analyses were used to assess the micropipette geometry and pore size. Electrochemical analysis involved filling and immersing the micropipettes with 1x PBS.

After selecting a region of interest, based on optical imaging, a biopsy can be performed by lowering the micropipette to the selected region by a predetermined depth (2μm) at a speed of 100μm/s whilst applying a negative voltage to prevent premature aspiration, which we optimally found to be −200mV. Our group has previously shown that micro and nanopipettes can be used to aspirate cytoplasmic content by taking advantage of electrowetting [31,46] and we investigated if this approach was suitable for the sampling from skeletal muscle fibres.

Figure 4 shows representative optical images (Figure 4a and b) and topographical scan (figure 4c), before (Figure 4a) and after (Figure 4b) the completion of the subcellular biopsy. The BF micrograph post biopsy (figure 4bi) shows what looks like an indentation in the tissue which is confirmed by the post biopsy a SICM scan. The indentation is about 3 μm deep and 7 μm wide as shown in the line profile in figure 4c and these dimensions are consistent with the micropipette size employed in this study (Figure). Interestingly, we did not observe any noticeable decrease in fluorescence post biopsy (Figure 4biii) indicating that a change in fluorescence cannot be used to assess the success of the biopsy as the fluorescent signal could originate from an area underneath the biopsy site.

**Figure 4.**
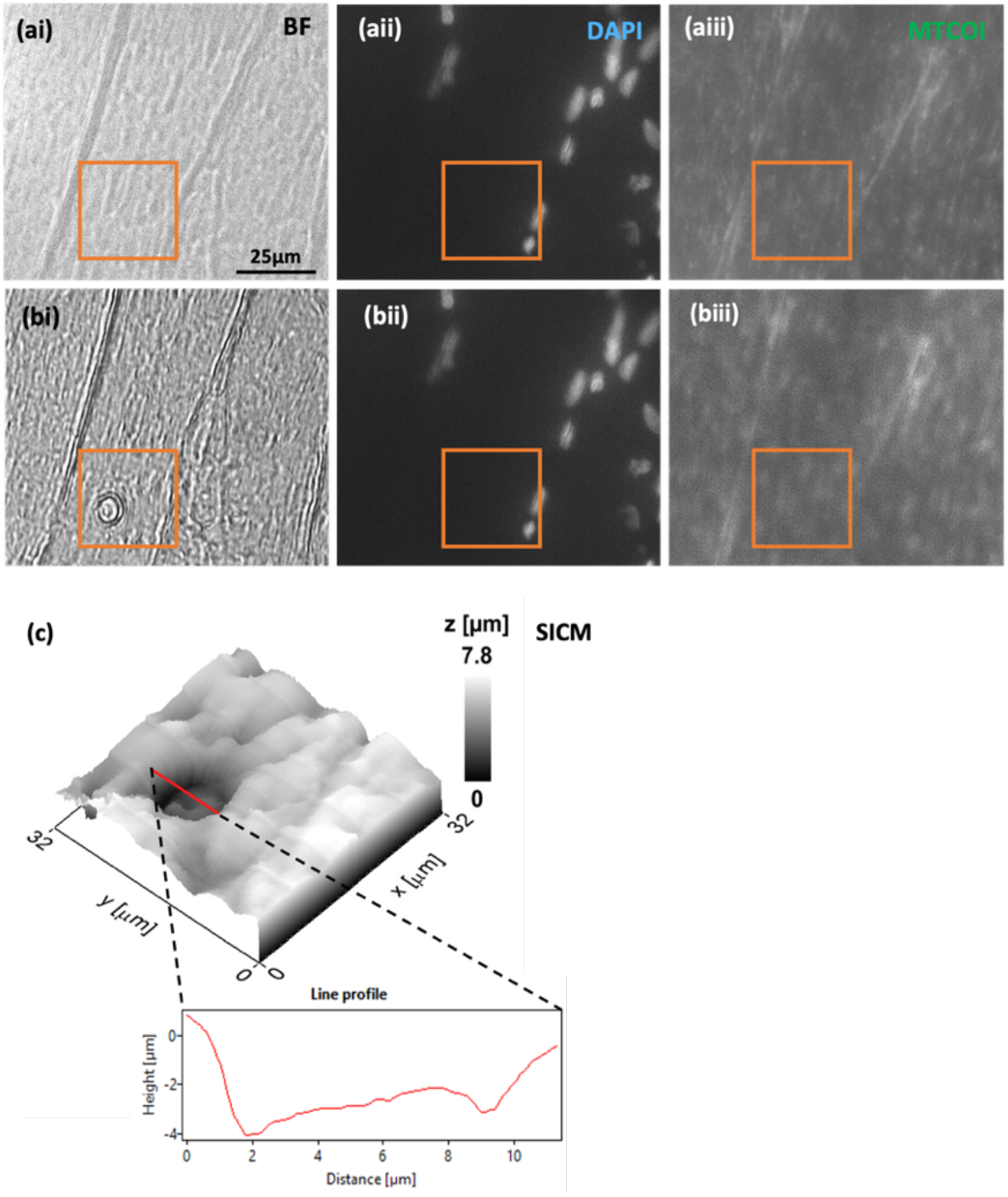
**(ai-aiii)** Pre subcellular biopsy images of skeletal muscle fibres, corresponding to BF **(ai)**, myonuclei stained with DAPI **(aii)** and IF staining of MTCOI **(aiii)**. **(bi-biii)** Post biopsy-images, again BF **(bi)**, DAPI **(bii)** and MTCOI IF staining **(biii)**. Evidence of successful biopsies is shown in BF **(bi)**, but a reduction in MTCOI fluorescence was not shown post-biopsy **(biii)**. The relative position of the biopsy to myonuclei is shown relative to DAPI staining, which indicates that biopsies taken are from the intermyofibrillar region and not from the perinuclear region. The biopsy sites in all images (pre- and post-biopsy) were delineated within an orange box. A topographical scan of the area shown within the orange box (32×32 μm) further supported the successful biopsy as shown by an indentation in the skeletal muscle fibre, measuring approx. 8μm in diameter, consistent with the biopsy site observed in BF **(c)**.

### Working range of subcellular mitochondrial biopsy

To compare the effectiveness of subcellular biopsy versus LCM, we performed the same triplex qPCR mtDNA assay on samples obtained with subcellular biopsy. In addition, we studied if the application of a negative voltage after the micropipette was inserted within the tissue was necessary to control the biopsy (Figure 5). The application of the negative voltage is necessary to control the electrowetting process but it does not seem to be necessary when a biopsy is performed within a tissue as we did not observe any statistically relevant difference when a biopsy was performed with electrowetting (negative voltage applied in tissue) and non-electrowetting (no voltage applied) (electrowetting mean CN/μl = 13.25; 95%CI= 2.87 - 24.22; non-electrowetting mean CN/μl = 13.55 95%CI= 5.76 - 20.74; Mann Whitney U, *p*>0.05, Figure 5). Although electrowetting appears to be critical to the isolation of mitochondria from cultured cells [29,31] we did not find any experimental evidence that is necessary to sample from a tissue slice.

**Figure 5.**
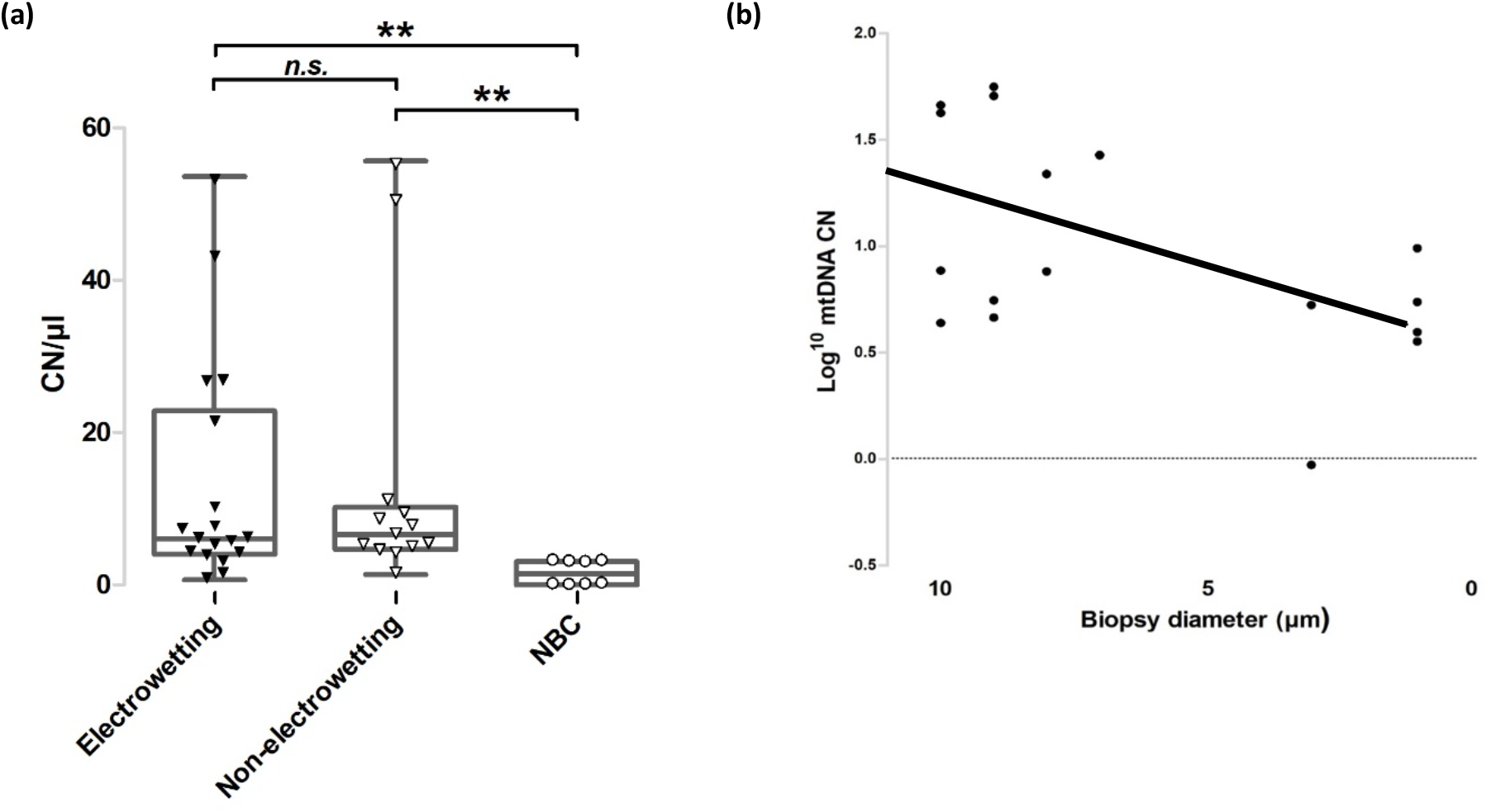
qPCR analysis of the biopsied samples. **(a)** Each data point corresponds a technical triplicate of each biopsy lysate sample. The box plot corresponds to the median and interquartile range of the data points. Data points shown as black triangles correspond to biopsies obtained with electrowetting (*n* = 11; mean = 13.25; median = 6.23) and points depicted as white triangles correspond to non-electrowetting (*n* = 7; mean = 13.55; median = 6.77). Points depicted as white dots correspond to non-biopsy controls (NBC; *n* = 5; mean = 1.71; median = 1.71). A significant difference in CN/μl was observed between biopsies acquired with (*p*<0.001, Mann-Whitney U test) and without electrowetting (p<0.001, Mann-Whitney U test) compared with non-biopsy controls (*p<* 0.03, unpaired t-test). There was no significant difference in CN/μl observed between biopsies acquired with electrowetting and non-electrowetting (*p*=0.76, Mann-Whitney U test). **(b)** The CN/μl of biopsies (log transformed, y-axis) was plotted against biopsy diameter as determined from topographical scans of biopsy site. The mean micropipette biopsy site diameter was 6.4μm. Biopsy site diameter and CN/μl were highly variably and a positive correlation that did no reach significance was observed between biopsy diameter and CN/μl (*n* = 11, *r* = 0.436, *p* >0.05).

Using subcellular biopsy, we were able to detect ~4 mtDNA CN from biopsies with a mean diameter of 6.4μm, based on SEM and SICM imaging data (n = 10; *sd* = 3.7, Figure 6, Supplementary Figures 1), Importantly, both electrowetting (Mann-Whitney U, p<0.005). and non-electrowetting Mann-Whitney U, p<0.005). MtDNA CN values were significantly higher than the non-biopsy controls (mean CN/μl = 1.71; 95%CI= 0.36-3.06).

Similar to LCM (Figure 2c), mtDNA CN was correlated to the biopsy diameter and whilst a statistically significant correlation was shown between total biopsy diameter and mtDNA CN (*n* = 10, *r* = 0.501, *p*<0.05, Figure 6), a correlation reaching statistical significance was not observed in either electrowetting (*n* = 6, *r* = 0.538, *p* = 0.087) or non-electrowetting (*n* = 4, *r* = 0.514, *p* = 0.238) biopsy groups.

## Discussion

Subcellular biopsy, as an adaptation of nanobiopsy, is a promising scanning probe technology for use in the isolation of organelles for downstream genomic analysis [26]. Whilst micromanipulation-based techniques and FFM have been previously utilized to acquire mitochondria from cells and tissues [35,36,47] and the use of scanning probes have been used to perform topographical scans of human tissue [48,49], up until now the utility of scanning probe technologies to aid the successful isolation of organelles from subcellular compartments within human tissue had yet to be demonstrated. We have optimized and utilized subcellular biopsy for the acquisition of mitochondria from skeletal muscle fibres for the first time, with superior control and sampling resolution offered by scanning probe technologies.

### Working range of Laser Capture Microdissection

A strong correlation was highlighted, in the qPCR data, between dissection size and mtDNA CN down to dissections of >20 μm in diameter, for dissection sizes <20μm in diameter a steep drop off in CN was observed. This is likely due to an unsuccessful dissection or due to ablative laser damage to mitochondria, mtDNA or even the muscle fibres themselves [50,51]. The latter is supported by observed scorching when areas <20 μm were circumscribed. This likely shows that the lower operative range of LCM, for isolating organelles and other biomolecules from human tissue, is likely to be in the region of 20μm. We have demonstrated that the subcellular biopsy has an operative range much lower than this. MtDNA CN data obtained from qPCR, coupled with topographical scans of the sampled biopsy sites, demonstrated a linear trend between CN and size, like that observed with LCM. There was no significant correlation between CN and biopsy size, this could be masked by a low *n* number and high variability in biopsy diameter.

### Optimization of subcellular biopsy in skeletal muscle fibres

The sampling of mitochondria, or other organelles, using scanning probe technologies had previously only been demonstrated in cultured cells [31,35]. Typically mitochondria from cultured cells tend to be smaller than in tissue [52,53] and so ensuring the micropipette pore size was sufficiently large to sample mitochondria was the first obstacle. The reported mean maximal diameter of subsarcolemmal (SS) mitochondria in human skeletal muscle tissue is 790nm, whilst the mean maximal diameter for intermyofibrillar (IMF) mitochondria is 1.2μm [54]. To avoid introducing sampling bias, micropipettes needed to be large enough to sample mitochondria from the upper range of any mitochondrial sub-population, whilst still being small enough to sample mitochondrial subpopulations unique to small distinct foci and to minimise the impact to the cellular environment of sampled skeletal muscle fibres [18,31].

### Subcellular biopsy allows the sampling of mitochondria from skeletal muscle fibres

Successful sampling of mitochondria from human tissues was confirmed by qPCR analysis, where more than two criterions were met. The limit of qPCR assay sensitivity was found to be 4.33 mtDNA CN/μl. The mean CN value of 13.25/μl was obtained with our cookie cutter approach. This is comparable with ~6-10 mtDNA copies per mitochondrion [55]. Compared with previous studies reporting isolation of a single mitochondrion [35,47], it is likely that this corresponds to several mitochondria but crucially our technology enables the sampling from larger or smaller regions of interest by simply adjusting the dimensions of the micropipette and it should theoretically allow the sampling of a single mitochondrion.

qPCR data suggests that electrowetting, whilst not necessarily detrimental, may not be necessary to perform biopsies in skeletal muscle tissue. This observation was exciting because it largely simplifies the operation of the SICM and negates the need for the organic solvent that needs to be used for electrowetting. A key benefit of subcellular biopsy is that the same platform employed for subcellular sampling can be used to acquire topographical scans of the region of interest before and after sampling [69]. Whilst, post-sampling topographical scans were useful in determining the success of sampling, real-time topographical scanning would improve the spatial resolution, speed and allows for automated sampling. Also, double or even multi-barrel micropipettes [46,56,58] could be used to allow for improved operations and enable multifunctional measurements. An exciting potential utility of subcellular biopsy, using SICM or other scanning probe systems, could be the acquisition and subsequent transplantation of mitochondria into cultured cells for investigative or therapeutic purposes [36,61]. The advantage of using an SICM system would be the ability to couple this with the topographical scanning capability of SICM [48,49].

## Conclusions

In this study, we demonstrated that a micropipette integrated within an SICM can successfully sample mitochondria from human skeletal muscle, or any human tissue, with a spatial resolution higher than the gold-standard LCM. We envision that this technology will enable the isolation and analysis of organelle populations from discrete foci within tissues for downstream molecular and structural analysis.

## Acknowledgments

This work was supported by a Wellcome Trust grant to the Wellcome Centre for Mitochondrial Research [203105]. A.B. acknowledges funding from the MRC DiMeN Doctoral Training Partnership [OSR/0200/2018/BURY]. A.V. acknowledges funding through a Sir Henry Wellcome Postdoctoral Fellowship [215888, https://doi.org/10.35802/215888] and F.M. and P.A. from the European Union’s Horizon 2020 research and innovation program under the Marie Skłodowska-Curie MSCA-ITN grant agreement no. 812398, through the single entity nanoelectrochemistry, SENTINEL.

